# Evidence of a Slower-Z effect in *Schistosoma japonicum*

**DOI:** 10.1101/2024.07.02.601697

**Authors:** Andrea Mrnjavac, Beatriz Vicoso

## Abstract

Sex-linked and autosomal loci experience different selective pressures and evolutionary dynamics. X (or Z) chromosomes are often hemizygous, as Y (or W) chromosomes often degenerate. Such hemizygous regions can be under greater efficacy of selection, as recessive mutations are immediately exposed to selection in the heterogametic sex (the so-called Faster-X or Faster-Z effect). However, in young non-recombining regions, Y/W chromosomes often have many functional genes, and many X/Z-linked loci are therefore diploid. The sheltering of recessive mutations on the X/Z by the Y/W homolog is expected to drive a Slower-X (Slower-Z) effect for diploid X/Z loci, i.e. a reduction in the efficacy of selection. While the Faster-X effect has been studied extensively, much less is known empirically about the evolutionary dynamics of diploid X or Z chromosomes. Here, we took advantage of published population genomic data in the female-heterogametic human parasite *Schistosoma japonicum* to characterize the gene content and diversity levels of the diploid and hemizygous regions of the Z chromosome. We used different metrics of selective pressures acting on genes to test for differences in the efficacy of selection in hemizygous and diploid Z regions, relative to autosomes. We found consistent patterns suggesting reduced Ne, and reduced efficacy of purifying selection, on both hemizygous and diploid Z regions. Moreover, relaxed selection was particularly pronounced for female-biased genes on the diploid Z, as predicted by Slower-Z theory.

## Introduction

Sex chromosomes, such as the X and Y of mammals, or the Z and W of birds, originate from standard pairs of autosomes. After they are coopted for sex determination, the two chromosomes typically stop recombining and start diverging from each other (Jay et al., 2024). This leads them to evolve differently from autosomes. The most striking aspect of this is the progressive degeneration of the non-recombining Y/W that is observed in many clades (Charlesworth, 2021). However, it has become increasingly appreciated that evolutionary rates on the X chromosome (or Z, but explained in terms of the X for simplicity) are also shaped by unusual evolutionary pressures. All else being equal, the effective population size of the X chromosome is ¾ the autosomal effective population size, while Y chromosomes have a population size of only ¼ the autosomal one (Vicoso and Charlesworth, 2009). Both X and Y chromosomes exhibit sex-biased transmission: the X resides in females ⅔ of the time, while the Y is in males 100% of the time (Furman et al, 2020). Furthermore, the degeneration of the Y chromosome (Bachtrog, 2013) leaves X-linked loci hemizygous in males. Selection is more efficient for hemizygous X-linked loci, where recessive mutations are always exposed to selection in males, compared to the autosomal loci. This should lead to higher rates of adaptive evolution on the X chromosome than autosomes if new beneficial are on average recessive, a hypothesis known as the Faster-X effect (Charlesworth et al., 1987, Vicoso and Charlesworth, 2006). Support for the Faster-X effect comes from the observation of elevated *dN/dS* on the X chromosome, or elevated values of α, the inferred proportion of nonsynonymous divergent sites that were fixed by positive selection (Meisel and Connallon, 2013), in various clades.

In particular, there is empirical evidence for high rates of non-synonymous evolution on the X in mammals and Drosophila, and on the Z in birds and arthropods (Meisel and Connallon, 2013, Charlesworth et al., 2018, Mank et al., 2010, Mongue et al., 2022). However, evidence suggests that this is not always driven by increased rates of adaptation. In particular, while there is evidence of increased rates of adaptive divergence on various X chromosomes (Garrigan et al. 2014, Avila et al. 2014, Veeramah et al. 2014, Kousathanas et al. 2014, Campos et al. 2014, Charlesworth et al. 2018), the Faster-Z effect has been interpreted as being the result of stronger drift on the Z chromosome of several species (Mank et al., 2010, Hayes et al., 2020, Chase et al., 2023, Mongue and Baird, 2024, but see Wanders et al., 2024). This is possibly due to the fact that the Z spends more time in males: males usually have a higher variance in reproductive success, resulting in the more extreme reduction in the effective population size for the Z chromosome than for the X (Vicoso and Charlesworth, 2009, Mank, Vicoso et al., 2010). In smaller populations, a higher proportion of mutations entering the population is effectively neutral, contributing to faster non-adaptive evolution (Mank et al., 2010, Mank, Vicoso et al., 2010). On the other hand, evidence of faster and more adaptive Z was found in some Lepidoptera, which typically have a larger population size than birds (which may make the Z chromosome less sensitive to the reduction in the effective population size) (Mongue et al., 2022, Villavicencio, 2024).

X-linked loci in young non-recombining regions, which still have a non-degenerated homologous region on the Y chromosome, are not hemizygous, but diploid in males, as they have a functional, albeit non-recombining, gametolog on the Y. Unlike loci on older, hemizygous X chromosomes, such “diploid X” loci are not expected to adapt faster than autosomal loci. New mutations that arise on a diploid X region are always heterozygous in males, and, if (partly) recessive, are (partially) sheltered from selection by the functional copy on the Y. This is expected to cause reduced efficiency of selection in males on diploid X region, slower adaptation of male-important genes and accumulation of deleterious mutations on male-important genes, i.e. a “Slower-X” effect (Mrnjavac et al., 2023).

The evolutionary patterns of young non-recombining regions on the X or Z have been studied less often, as population data is needed to detect very young non-recombining regions with non-degenerated Y counterparts (Darolti et al., 2022), but a few have found some support for Slower-X/Z evolution. Neo-X regions (with the corresponding Y chromosomes showing intermediate levels of degeneration) in several Drosophila species experience accelerated pseudogenization, driven by the loss of male-important genes (Nozawa et al., 2016, Nozawa et al., 2021). In the plant *Silene latifolia* there is evidence of relaxed purifying selection on young X-linked genes with a non-degenerated Y homolog (Krasovec et al., 2018). Recently, a study in the butterfly genus Leptidea provided direct empirical evidence of reduced efficiency of selection for female-biased and unbiased genes on the young non-recombining region of the Z chromosome with a non-degenerated W, i.e. of a Slower-X (Slower-Z) effect (Hook et al., 2024). On the other hand, some studies have found similar rates of divergence for diploid X/Z genes as for (pseudo)autosomal genes. The young X-linked region of the plant *Salix dunni* is enriched for transposable elements and pseudogenes, but divergence of X-linked genes is similar to the autosomal divergence, possibly because the X-linked region is very young and there was no time for non-adaptive substitutions to accumulate (He et al., 2021). Similarly, in Sylvioidea songbirds, there is no difference in evolutionary rates between the neo-Z and autosomes (Leroy et al., 2021). Darolti et al. (2023) further showed that while Faster-X correlates with hemizygosity in various species of poeciliid fishes, no evidence of increased drift or differences in divergence rates could be detected between diploid X chromosomes and their respective autosomes. Therefore, the broad relevance of the Slower-X effect in taxa with young sex-linked regions is still to be fully explored.

Blood flukes (genus Schistosoma) are a promising model for studying the evolutionary dynamics of sex-linked regions of different ages. While they all share an ancestral pair of ZW chromosomes, the non-recombining part of the sex chromosomes has been expanded independently in different lineages (Picard et al., 2018). A very young non-recombining region of the Z chromosome has been recently identified in the Asian species *Schistosoma japonicum* (Elkrewi et al. 2021, Xu et al., 2023). This region has over 700 functional W genes (ZW dS < 0.085), which makes the corresponding Z region diploid but non-recombining in females. We expect such a region to be under reduced efficiency of selection in females, compared to autosomes and hemizygous Z (Mrnjavac et al., 2023). Here, we use publicly available comparative (Protasio et al., 2012, Luo et al., 2022), population (Luo et al., 2022) and expression data (Wang et al., 2017) to test those predictions.

## Methods

### Strata determination

To identify hemizygous and diploid Z regions, we performed female-to-male coverage analysis and male-to-female Fst analysis as in Elkrewi et al. (2021), using the recently published male *S.japonicum* genome assembly (GCA_021461655.1) (Luo et al., 2022). Briefly, female (SRR6841388) and male (SRR6841389) *S.japonicum* reads were separately mapped to the *S.japonicum* male genome using *bowtie2* (Langmead and Salzberg, 2012). Only uniquely mapped reads were kept. Coverage for male and female reads was calculated with *soap.coverage* (Luo et al. 2012) per 10000 bp windows. Log2(F/M coverage) was calculated and visualised in R (R Core Team, 2023). Coordinates of the hemizygous Z region were determined as the limits of the Z chromosome region where log2(F/M coverage) values are centred at −1, meaning there are twice as many reads in males compared to females (Z chromosome coordinates: 24470001-49640001).

The Fst analysis also followed the approach of Elkrewi et al. (2021), but using the new chromosome-level genome assembly. Fst between male and female reads (PRJNA650045, sex of the individual library was determined from Elkrewi et al., 2021) was calculated with *vcftools* (Danecek et al., 2011) and visualised in R. The diploid Z region was determined as the region for which the male:female Fst values were consistently above the 95 percentile of the distribution across the genome (Z chromosome coordinates: 49640001-76240000). In this region 62.67% of windows had male:female Fst values above the 95 percentile of the genome-wide distribution and 90.42% of reads had male:female Fst values above the 90 percentile of the genome-wide distribution.

### Sex-biased expression analysis

Publicly available whole-body expression data was downloaded for 24 male and 24 female *S.japonicum* individuals, (PRJNA343582, Wang et al., 2017). Normalised gene expression was obtained per gene, per sample, using *kallisto* (Bray et al., 2016) and *sleuth* (Pimentel et al., 2017). We filtered out the genes with no expression in both of the sexes. We estimated sex-biased gene expression as SPM (Kryuchkova-Mostacci and Robinson-Rechavi, 2017), the square of mean expression in females divided by the sum of the square of mean expression in females and the square of mean expression in males, using R. SPM=0 corresponds to male-limited expression, while SPM=1 corresponds to female-limited expression. For further analyses, genes with SPM values lower than 0.3 were assigned as male-biased, and genes with SPM values above 0.7 were assigned as female-biased. Distributions of SPM values in autosomes, hemizygous Z and diploid Z regions were visualised and compared in R. Sex-bias distribution was compared between the Z chromosome and autosomes with Kolmogorov–Smirnov test and Mann-Whitney-Wilcoxon test.

### Divergence inference

We identified orthologs between *S.japonicum* and closely related species *S.mansoni* (65% median synonymous divergence, Picard et al., 2018) as the best reciprocal blat (Kent, 2002) hits between *S.japonicum* and *S.mansoni* coding sequences (assembly version GCF_000237925.1, Protasio et al., 2012) (we chose the longest coding sequence per gene for the analysis). Orthologs were aligned using TranslatorX with the “gblocks” option (Abascal et al., 2010). Divergence between orthologs was calculated with KaKs_Calculator 2.0 (Wang et al., 2010). Yang-Nielsen estimates of Ka/Ks were obtained per gene, as well as the number of nonsynonymous and synonymous substitutions per gene. These parameters were visualised and compared in R.

### Polymorphism inference

We downloaded a publicly available population genomic dataset (PRJNA789681), including whole genome sequences of 48 *S.japonicum* adult male individuals sampled from several locations in South-East Asia (we did not include the Taiwan and the Philippines subpopulations in our analysis as those subpopulations have extremely reduced levels of diversity and could have biased our analysis), corresponding to their worldwide range (Luo et al., 2022). We trimmed the reads with *Trimmomatic* (Bolger et al., 2014). Trimmed paired reads were mapped to the *S.japonicum* male genome assembly (GCA_021461655.1) using *bowtie2* with *--end-to-end* and *--sensitive* parameters, separately for every individual. Non-uniquely mapped reads were removed. SAM files were reformatted into sorted BAM files using samtools (Li et al., 2009). Variant calling was performed with *bcftools mpileup* option (Li et al., 2009), using (48) 72 individual bam files as input. Variants were filtered by quality, *bcftools view -i ‘%QUAL>=20’*, only biallelic sites were kept, *--max-alleles 2*, and indels were removed, *--exclude-types indels*. bcf file was reformatted into vcf file. Rare variants (maf<15%) were removed with *vcftools* using *--maf 0.15 --max-missing 0.9* options. Polymorphic sites were annotated as synonymous or nonsynonymous using *snpEff* and *SnpSift* (Cingolani et al., 2012).

### Population genomic analyses

#### α

α per gene was calculated as 1-((number of nonsynonymous polymorphisms per gene (Pn) / number of synonymous polymorphisms per gene (Ps))/(number of nonsynonymous substitutions per gene (Dn) / number of synonymous substitutions per gene (Ds))) (Smith and Eyre-Walker, 2002, Charlesworth and Charlesworth, 2010), after removing variants below 15% frequency (Fay et al., 2001, Al-Saffar and Hahn, 2022), using R. Distributions of α values for different categories of sex-bias, and different genomic regions: hemizygous Z, diploid Z and autosomal one, were visualised and statistically compared in R.

#### Direction of Selection

In addition to α, we calculated Direction of Selection (DoS) as DoS=Dn/(Dn+Ds)-Pn/(Pn+Ps) (Stoletzki and Eyre-Walker, 2011).

#### Nucleotide diversity

Nucleotide diversity along the genome, in 10000 bp windows, was calculated using pixy (Korunes and Samuk, 2021).

## Results

### Hemizygous and diploid regions of the Z chromosome

Elkrewi et al. (2021) and Xu et al. (2023) recently described evolutionary strata of different ages along the Z chromosome of *S. japonicum*, and in particular the presence of a large section of the ZW pair that no longer recombines, but still exists on the W. However, a highly fragmented genome was used in Elkrewi et al. (2021), and no population genomics data was used to infer young non-recombining regions in Xu et al. (2023). We therefore set out to define precise boundaries of the diploid and hemizygous Z regions on the published chromosome-level assembly of *S. japonicum* (Luo et al., 2022). Using both coverage patterns and genetic differentiation between a population of males and females, we recovered large contiguous hemizygous and non-recombining but diploid Z regions (Xu et al., 2023, Elkrewi et al. 2021) (**Figure 1**). In particular, a large region where female coverage is consistently half of male coverage is consistent with the degeneration of the homologous region of the W chromosome, i.e. this Z chromosome region is haploid or hemizygous in females. A second region shows no difference in coverage between males and females, but shows a high level of genetic differentiation between the Z and W, measured as male-to-female Fst, consistent with a recent loss of recombination between the Z and W, and a non-degenerated homologous region on the W chromosome. The hemizygous and diploid Z regions contain, respectively, 703 and 624 genes.

**Figure 1.**
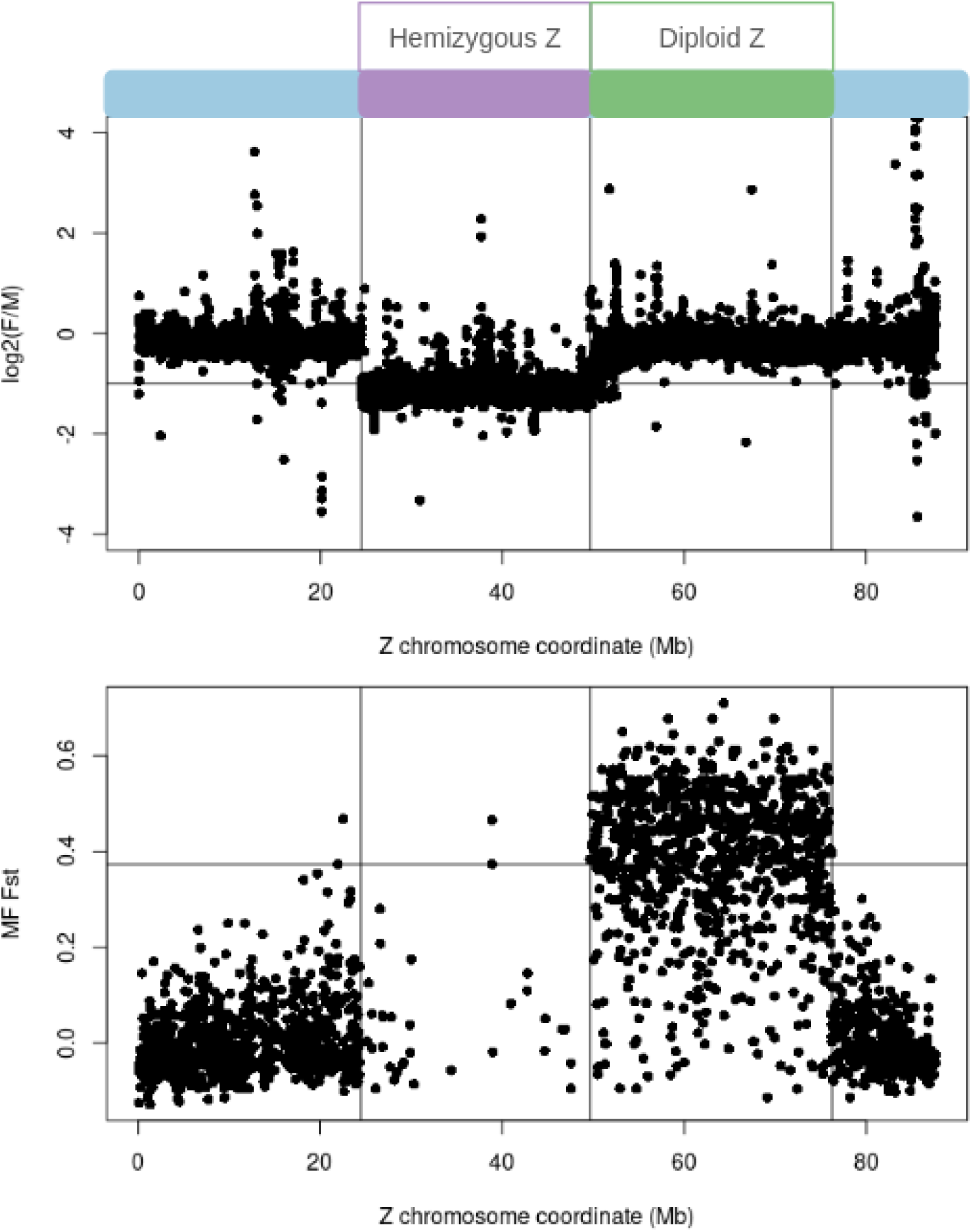
*Schistosoma japonicum* Z chromosome strata: hemizygous and diploid Z regions. Upper panel: Log2(Female coverage / Male coverage) along the Z chromosome. Hemizygous Z region has male coverage that is two times the female coverage. Lowe panel: Male-to-Female Fst along the Z chromosome. Diploid Z region has equal coverage in males and females,but has high levels of Male-to-Female Fst.

### Lower effective population size on the Z chromosome

Since males have two Z chromosomes but females only have one (compared to two sets of autosomes in each sex), the expected effective population size of the Z is ¾ of that of the autosomes. We estimated total genetic pairwise diversity (π) from a population of 48 males from 6 sampling locations, and used it to infer the effective population size of the hemizygous and diploid Z regions relative to that of the autosomes in *Schistosoma japonicum*. Both hemizygous and diploid Z regions show lower than expected nucleotide diversity compared to autosomes, with the Z/A ratio of median nucleotide diversity 0.37 for diploid Z region and 0.09 for hemizygous Z region (**S.Figure 1**), which suggests that effective population size of the Z chromosome could be even lower than ¾ of the autosomal effective population size. Similar estimates were obtained when only synonymous sites were used to calculate diversity (Z:A ratios of 0.395 and 0.235 for the diploid and hemizygous regions). Given the apparent young age of the diploid Z region, we took advantage of the reduced π on the diploid Z to check that the loss of ZW recombination was found in every population. Two populations had extremely reduced levels of diversity and were excluded from further analysis. In each of the other populations, the diploid Z region had reduced levels of diversity compared to the autosomes (p<2*10^-16^, Mann-Whitney-Wilcoxon test), confirming that it is non-recombining throughout the geographical range of the species (**S.Figure 1B**). Such a reduction was not observed in the pseudoautosomal region (p>0.1).

### Reduced efficacy of purifying selection on both the hemizygous and diploid Z regions

We first measured the divergence between *S.japonicum* and the closely related species *S.mansoni* to estimate synonymous (Ks) and nonsynonymous (Ka) substitution rates per gene. **Figure 2** shows the distribution of Ka/Ks for hemizygous Z, diploid Z and autosomes: Ka/Ks is significantly higher on the hemizygous Z compared to both autosomes and to the diploid Z (W = 1178163, p-value = 1.327*10^-13^, W = 119932, p-value = 0.0003441, respectively), while diploid Z genes show a slight increase compared to the autosomes (W = 1243012, p-value = 0.02237). Synonymous divergence is significantly lower on the Z chromosome, with hemizygous Z exhibiting the lowest synonymous divergence (hZ vs A: W = 1918300, p-value < 2.2*10^-16^, dZ vs A: W = 1565712, p-value = 1.656*10^-11^, hZ vs dZ: W = 158188, p-value = 2.493*10^-5^). Overall these results support the faster protein divergence of Z-linked genes compared to the autosomes.

**Figure 2.**
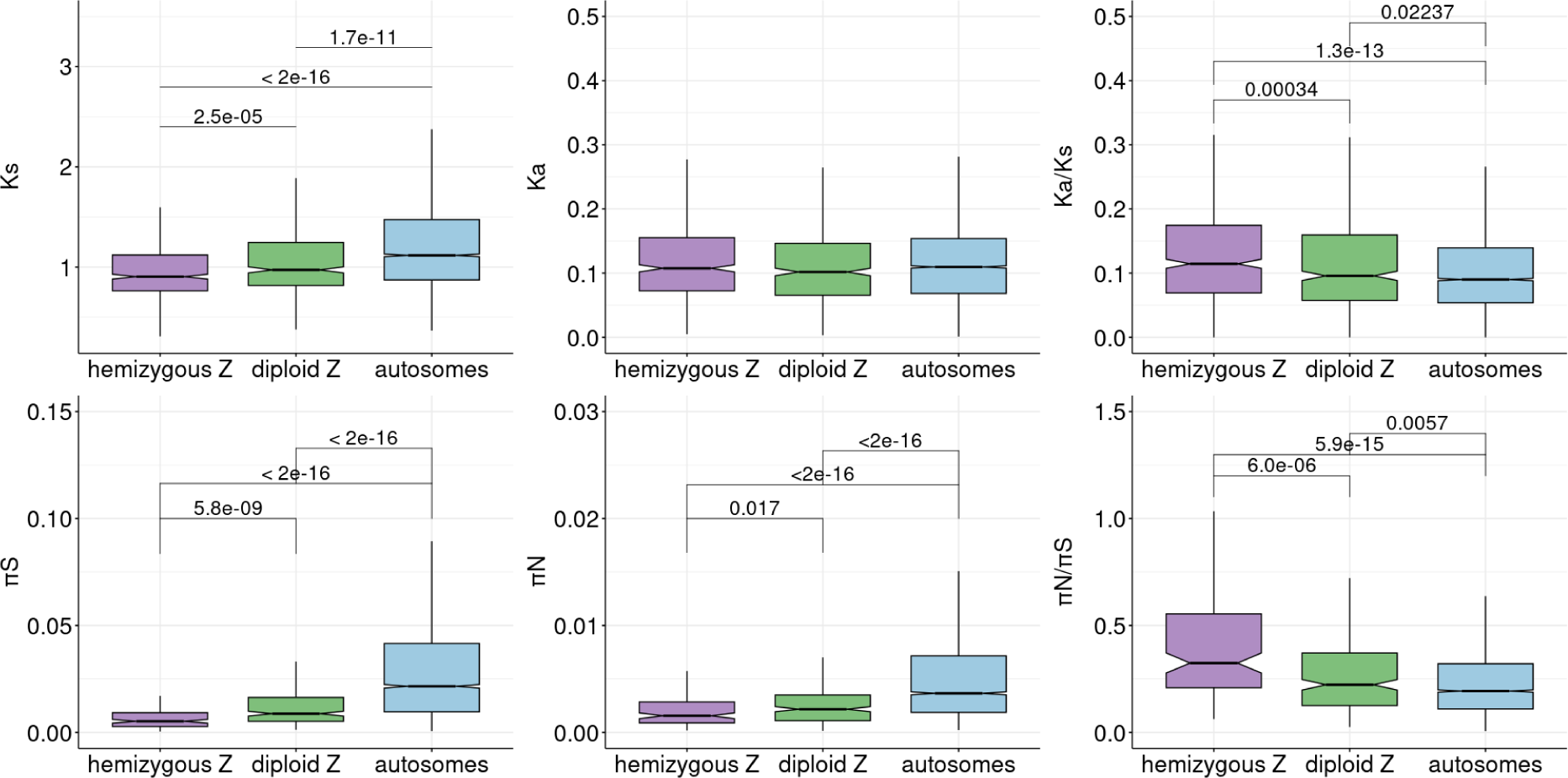
Synonymous (Ks) and nonsynonymous divergence (Ka), and their ratio (Ka/Ks), nonsynonymous (πS) and synonymous diversity (πN) and their ratio (πN/πS) for hemizygous and diploid Z and autosomes.

In order to investigate whether this fast evolution of Z-linked genes was driven by an increase in positive selection or by a decrease in the efficacy of purifying selection, we obtained estimates of synonymous and nonsynonymous polymorphism across the sampled populations (excluding the two that did not harbour any diversity). The Z chromosome has higher levels of nonsynonymous to synonymous diversity compared to the autosomes (hZ vs A: p=5.9*10^-15^, dZ vs A: p= 0.0057) (**Figure 2**). This is in line with the reduced effective population size and the resulting reduced efficiency of selection in removing slightly deleterious mutations from the population. We also calculated α per gene, a regularly used measure of adaptive evolution based on the McDonald-Kreitman test (McDonald and Kreitman, 1991, Smith and Eyre-Walker, 2002, Charlesworth and Charlesworth, 2010). Positive α values suggest positive selection, while negative α values mean there is an excess of nonsynonymous polymorphisms segregating in the population. This excess is usually caused by segregating slightly deleterious mutations, that is, lower efficiency of selection, or, balancing selection (Charlesworth and Charlesworth, 2010). We removed rare polymorphic sites (with minor allele frequency below 15%) from the analysis to minimize the contribution of deleterious mutations segregating at low frequencies. We also removed the genes with no polymorphism for the downstream analysis, which greatly reduced the number of genes: in the hemizygous Z region up to 80% of the genes did not exhibit any polymorphism after filtering out rare variants, while in the diploid Z region and autosomes, from 20% to 60% of genes exhibited no polymorphisms. This reflects extremely low levels of nonsynonymous and synonymous polymorphisms segregating on the hemizygous Z region (**Figure 2**), congruent with the extreme reduction in the population size for hemizygous Z compared to the rest of the genome (**S.Figure 1**). The small number of genes in some categories, especially on hemizygous Z, greatly reduced our statistical power. It should be noted that our values of α likely underestimate the true proportion of nonsynonymous substitutions fixed by positive selection, however, here we are interested in relative differences in the strength of selection in different genomic regions. Furthermore, α values for diploid Z region should be interpreted taking into account that observed higher diversity for female-biased genes is not reflected in higher divergence, as this region only recently became diploid and its divergence reflects autosomal patterns. In agreement with purifying selection being relaxed on both the hemizygous and diploid Z, genes in both regions showed reduced α values compared with autosomal genes (p=4.5*10^-10^ and p=0.005) (**S.Figure 3A**). Once again, the effect was stronger for the hemizygous Z region than for the diploid Z.

In addition to α, we calculated a second metric of selection strength, the Direction of Selection (DoS) (Stoletzki and Eyre-Walker, 2011), and the results were qualitatively similar (**S.Figure 4A**). The lower values of DoS observed for both hemizygous and diploid Z genes compared to autosomal genes (p=4.7*10^-10^ and p=0.0043) suggest that there is an excess of nonsynonymous polymorphisms that reach high frequencies in the population (as we removed rare variants) on both regions of the Z.

### Different evolutionary dynamics of hemizygous and diploid Z-linked sex-biased genes

Genes with sex-specific functions are expected to evolve differently on hemizygous and diploid Z-linked regions. To test this hypothesis, we used sex-specific patterns of expression as a proxy for function. We measured sex bias in the expression of the *S.japonicum* whole body throughout the development, as SPM (specificity metric, Kryuchkova-Mostacci and Robinson-Rechavi, 2017), where 0 corresponds to male-specific expression and 1 corresponds to female-specific expression. **S.Figure 2**. shows distributions of sex-biased expression on autosomes, in the diploid Z region and in the hemizygous Z region. The hemizygous Z region is significantly masculinized (D = 0.54589, p-value < 2.2*10^-16^, W = 3170339, p-value < 2.2*10^-16^), in agreement with its incomplete mechanism of dosage compensation (Picard et al., 2018), while the diploid Z region exhibits a small shift towards male-biased expression (D = 0.11372, p-value = 2.487*10^-6^, W = 1953576, p-value = 0.0003627).

**Figure 3.** shows nonsynonymous to synonymous substitution rates and genetic diversity as a function of sex-bias and genomic region: hemizygous Z, diploid Z and autosomes. Unbiased genes generally follow the trends described above for all genes: both hemizygous Z and diploid Z genes have increased Ka/Ks (though only significantly so in the case of diploid genes, p=0.025) and increased πN/πS compared with autosomal genes (p=9.7*10^-7^ and p=0.027 for hemizygous and diploid Z genes), consistent with reduced efficacy of selection on both parts of the Z. This is also supported by their reduced α values compared with autosomal values (**S.Figure 3B**, p=0.00038 and p=0.00781 for hemizygous and diploid Z genes).

**Figure 3.**
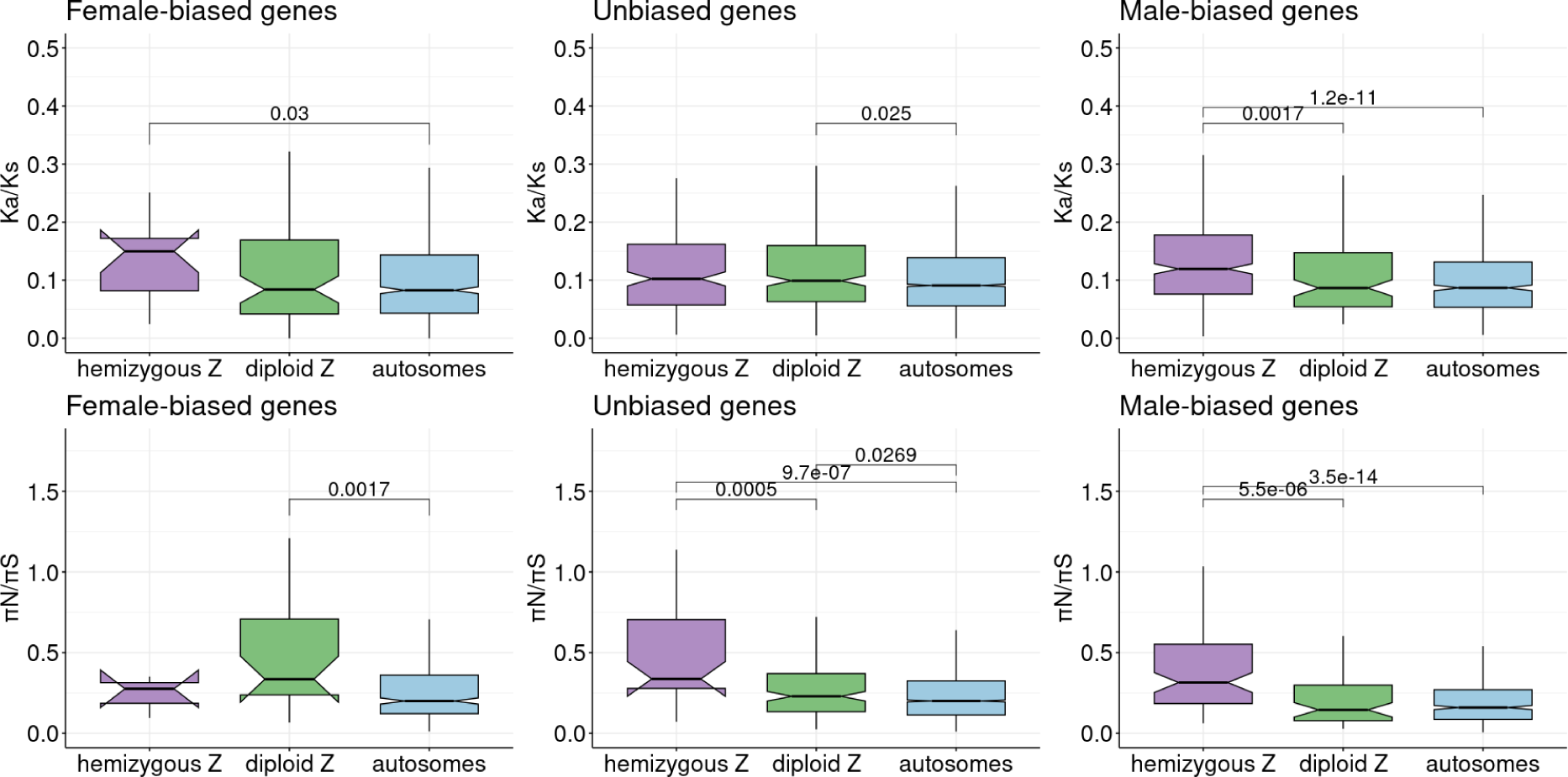
Ka/Ks and πN/πS for hemizygous and diploid Z and autosomes as a function of sex-bias.

In the hemizygous Z region, a key prediction is that genes that function primarily in females are expected to be under stronger efficacy of selection than equivalent autosomal genes, potentially leading to higher rates of adaptive divergence. The ratio of nonsynonymous to synonymous divergence (Ka/Ks) for female-biased genes on the hemizygous Z is significantly higher than for autosomal female-biased genes (W = 7432, p-value = 0.03918). Female-biased genes on the hemizygous Z also have a higher median Ka/Ks than female-biased genes on the diploid Z, however, the difference is not significant (W = 718, p-value = 0.1146), possibly due to low statistical power, as there are only 15 and 76 genes, respectively, in these categories. πN/πS values did not differ between female-biased genes on the hemizygous Z and on the autosomes, and α trended towards higher values for the former (though not significantly so), suggesting that positive selection acting on hemizygous mutations may contribute to the observed increase in protein coding divergence. Male-biased genes on the hemizygous Z also showed higher Ka/Ks than male-biased genes on autosomes (W = 137936, p-value = 8.587*10^-12^) but this was in this case associated with elevated levels of πN/πS (hZ vs A mb: W = 35139, p-value = 3.512*10^-14^) and reduced values of α (p=2.1*10^-15^), consistent with a primary role of relaxed purifying selection.

On the diploid Z, the expectation is that female-biased genes should be under strongly reduced efficacy of selection. Neither female-biased nor male-biased genes on the hemizygous Z showed a significant difference in Ka/Ks when compared to their respective autosomal controls (W = 29213, p-value = 0.8083, W = 39549, p-value = 0.239, respectively). While πN/πS did not differ between diploid Z and autosomal male-biased genes, suggesting the two are under similar selective pressures, female-biased genes on the diploid Z had higher πN/πS and lower α than their autosomal counterparts (dZ vs A fb: p = 0.001672 and p=0.0078). (**Figure 3, S.Figure 3B**). This is generally in line with our predictions that mutations can freely accumulate on female-biased genes in diploid Z, as they are sheltered from selection by the functional gemetolog on the W.

## Discussion

The sex chromosomes of *Schistosoma japonicum*, with their clearly distinguishable hemizygous Z and diploid Z regions, that is, strata with degenerated W and with non-degenerated W (Elkrewi et al., 2021, Xu et al., 2023), provide an ideal opportunity to study the evolutionary dynamics of young and old sex-linked regions in the same species. Our results suggest that the effective population size (Ne) of the Z chromosome in *S. japonicum* is much lower than the expected ¾ of the autosomal effective population size. Since neutral diversity levels are a function of Ne times the mutation rate μ, it is possible that the difference is driven by differences in μ. Indeed, the lower synonymous divergence (Ks) observed for the Z compared with the autosomes suggests that mutation rates may be lower on the Z. To account for this, we also compared the distribution of π/Ks, which should control for the mutation rate. Values for the Z were still lower than ¾ of those of the autosomes (hZ/A=0.2584, dZ/A=0.4290, S.Figure 5), suggesting a true reduction in Ne. The effective population size of Z chromosomes is expected to typically be smaller than the effective population size of X chromosomes, because Z chromosomes spend most of the time in males, and males often have a larger variance of reproductive success, which decreases their effective population size (Caballero, 1995, Charlesworth, 2001, Laporte and Charlesworth, 2002, Vicoso and Charlesworth, 2009, Mank, Vicoso et al., 2010). While the variance in reproductive success of males and females of *S. japonicum* is not known, adult populations of adults are typically male-biased (Beltran and Boissier, 2010). Given the largely monogamous reproductive mode of schistosome parasites (Beltran and Boissier, 2008), this may lead to a substantial proportion of males remaining unpaired, thereby increasing the variance in their reproductive success.

Interestingly, the Ne of the hemizygous Z is lower than the Ne of the diploid Z. One possibility to explain this is that the hemizygous Z region has been non-recombining for a longer amount of time: if loss of recombination with the W occurred very recently, the diploid Z may still not have lost all the standing variation that it harbored when it was a pseudoautosomal region. This is however unlikely to fully explain the pattern, as the reduction in Ne following a decrease in population size should occur fairly rapidly (as the long-term Ne is simply the harmonic mean of the population sizes over generations, Kalinowski and Waples, 2002). The lower Ne observed on the Z chromosome could also be due to the stronger effect of linked selection. The effect of linked selection should be particularly strong on the hemizygous Z, due to recessive mutations being exposed to selection. While we did not detect evidence of stronger positive selection on the hemizygous Z than on the autosomes, a recent study did detect a few loci under strong selection (Zhou et al., 2024), which may have contributed to reducing its genetic diversity. Additionally, it is possible that the hemizygous Z region does not recombine even in males (or has very low recombination rates), which would further reduce diversity. While no linkage map is available for *S. japonicum*, the Z-specific region of its close relative *S. mansoni*, which is partly shared with *S. japonicum*, has normal levels of recombination in males (Criscione et al., 2009). It therefore seems likely that a combination of factors drives the strong reduction in Ne that we observe.

Consistent with this reduced effective population size, our results suggest that the evolution of the Z chromosome in *S.japonicum* is dominated by the effect of relaxed purifying selection. This is in line with the general pattern of faster rates of evolution on the Z chromosome, which are often caused by drift due to its smaller effective population size (Mank, Vicoso et al., 2010, Mank et al., 2010, Hayes et al, 2020, Chase et al., 2023, Mongue and Baird, 2024). Although both the hemizygous and diploid Z regions are under reduced efficacy of purifying selection, we could to some extent test the differential expectations of the “faster-Z” and “slower-Z” effects by focusing on sex-biased genes. The effect of drift is expected to be counteracted to some extent by strong haploid selection for female-biased genes on hemizygous Z (Vicoso and Charlesworth, 2006). Consistent with this, female-biased genes located on the hemizygous Z region had a slightly increased Ka/Ks, but not πN/πS, when compared to autosomal female-biased genes. However, since only a handful of genes had sufficient polymorphism in our dataset, we could not obtain a significant signal of Faster-Z effect when using α inferences. On the other hand, in addition to reduced Ne, in the young diploid Z region, selective constraints should be relaxed due to the sheltering effect of functional gametologs on the W, and this effect should be stronger for genes expressed primarily in females. The effect of sheltering is supported by the fact that female-biased genes have the highest πN/πS, and the lowest inferred α, of the genes in the diploid Z region. Taken together, these results confirm that diploid and hemizygous sex-linked regions have different evolutionary dynamics, and that genes that function predominantly in one sex are primarily affected (assuming that sex-biased gene expression is a good proxy for sex-biased function).

Several theoretical models predict that Z-chromosomes may become “masculinized” over time, i.e. they may lose genes with female-specific functions and gain genes that work primarily in males (Gurbich and Bachtrog, 2008, Mrnjavac et al., 2023). An excess of Z-linked genes of *S. japonicum* are indeed male-biased in their expression. In the hemizygous Z region, masculinized expression can to a large extent be explained by the incomplete dosage compensation system found in this group: the Z chromosome is upregulated in both sexes, and has higher expression in males, since males have two copies of the Z (Picard et al., 2018). Whether an ancestral enrichment in genes with male-specific functions favored the evolution of such an unusual regulatory mechanism has yet to be tested. In the diploid Z region, expression patterns are more similar to autosomal ones, as we are capturing expression from the W gametologs in females, but a significant bias towards higher male expression was observed. This could have two (non-mutually exclusive) explanations: 1. Genes on the W may have undergone some regulatory degeneration, leading to their lower expression; 2. Genes on the Z may have become masculinized, ie male-beneficial mutations may have favored their increased expression in males and/or decreased expression in females, as predicted by the Slower-Z hypothesis. A recent study found similar expression levels from the W and the Z in the diploid Z region (Elkrewi et al., 2021); however, there was very limited power as only a small subset of genes were sampled. Future work comparing the expression of the Z and the W over the whole region, as well as patterns of expression of these genes in species where they are not sex-linked, may shed light on which of these hypotheses is driving this shift.

Our study illustrates different evolutionary dynamics of old and young sex-linked regions. Together with other studies on young sex-linked regions in butterflies of genus Leptidea (Hook et al.,2023), plant *Silene latifolia* (Krasovec et al., 2018), and several Drosophila species (Nozawa et al., 2016, Nozawa et al., 2021), our study suggests that Slower-X (or Slower-Z) effect might be widespread in young sex-linked regions. This body of work also illustrates the importance of studying non-model species, where diploid Z and X regions might be common, but underreported, as well as using population data for studying ongoing evolutionary processes.

