## Supplementary figures for "Evidence of a Slower-Z effect in *Schistosoma japonicum*"

A)

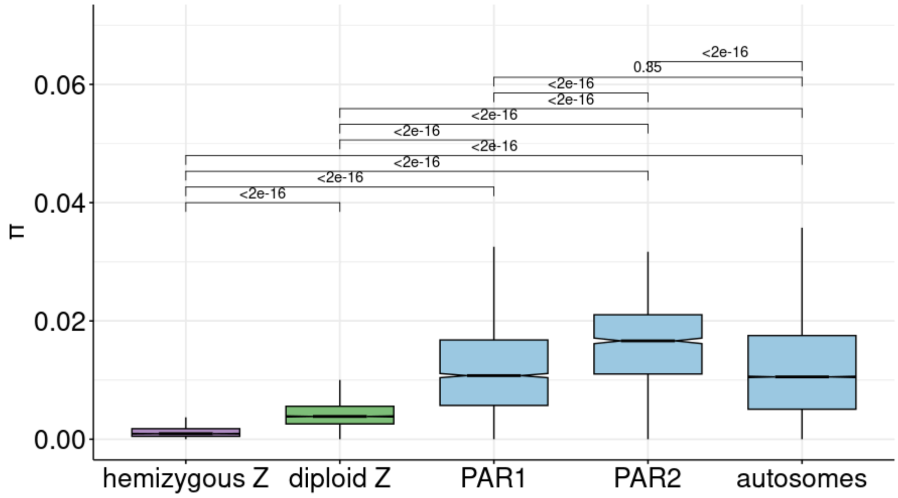


B)

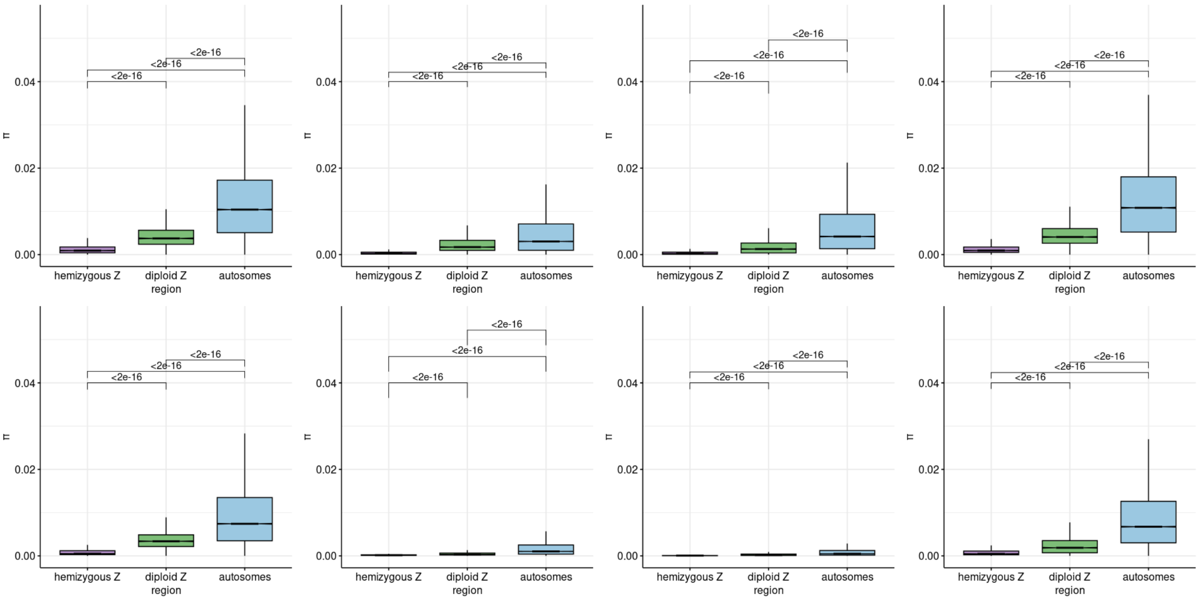


**S.Figure 1.** A) Nucleotide diversity (π) of hemizygous Z, diploid Z and pseudoautosomal regions and autosomes. B) nucleotide diversity in subpopulations.

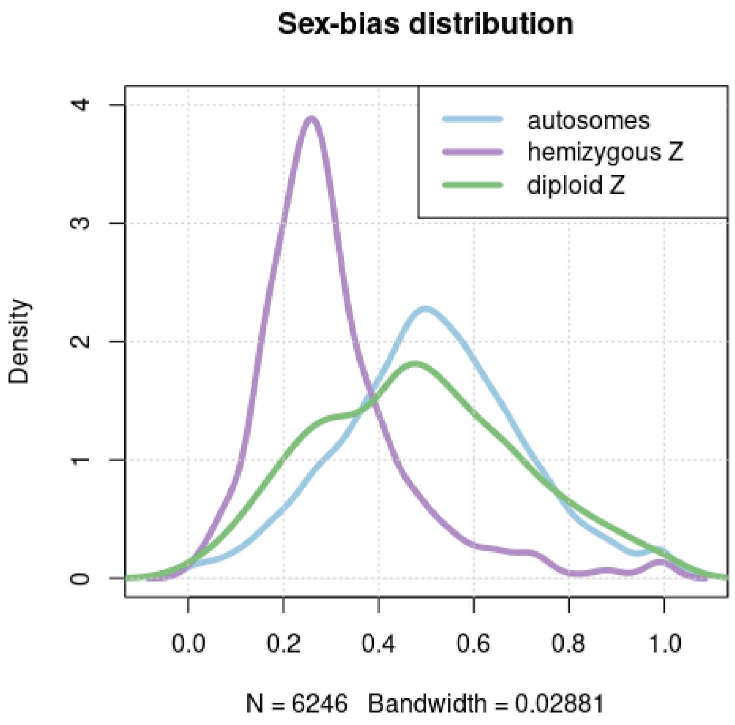


**S. Figure 2.** Sex-bias distribution in autosomes, diploid Z region and hemizygous Z region.

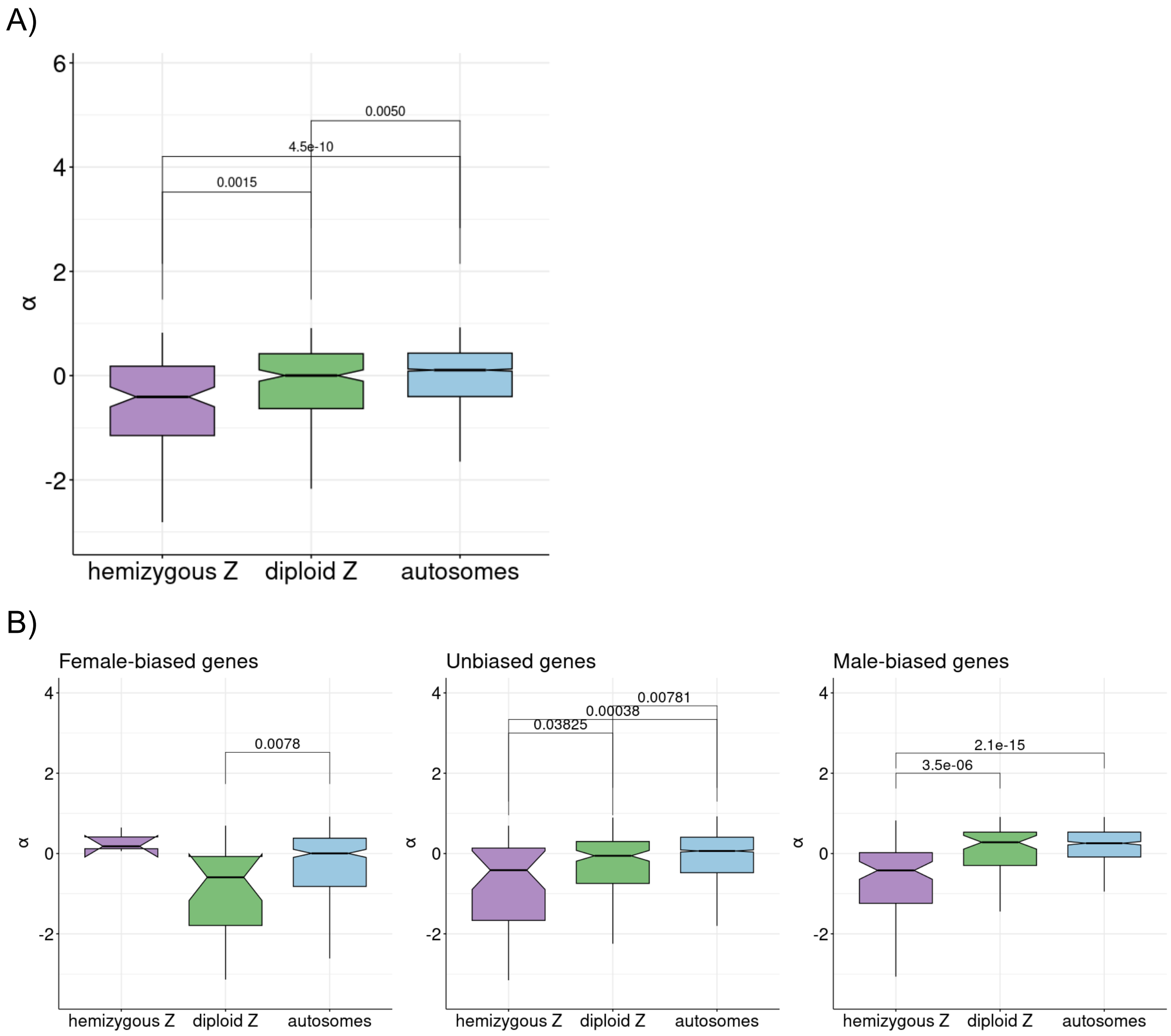


**S.Figure 3. A** α as a function of genomic location **B** α as a function of sex-bias and a genomic location.

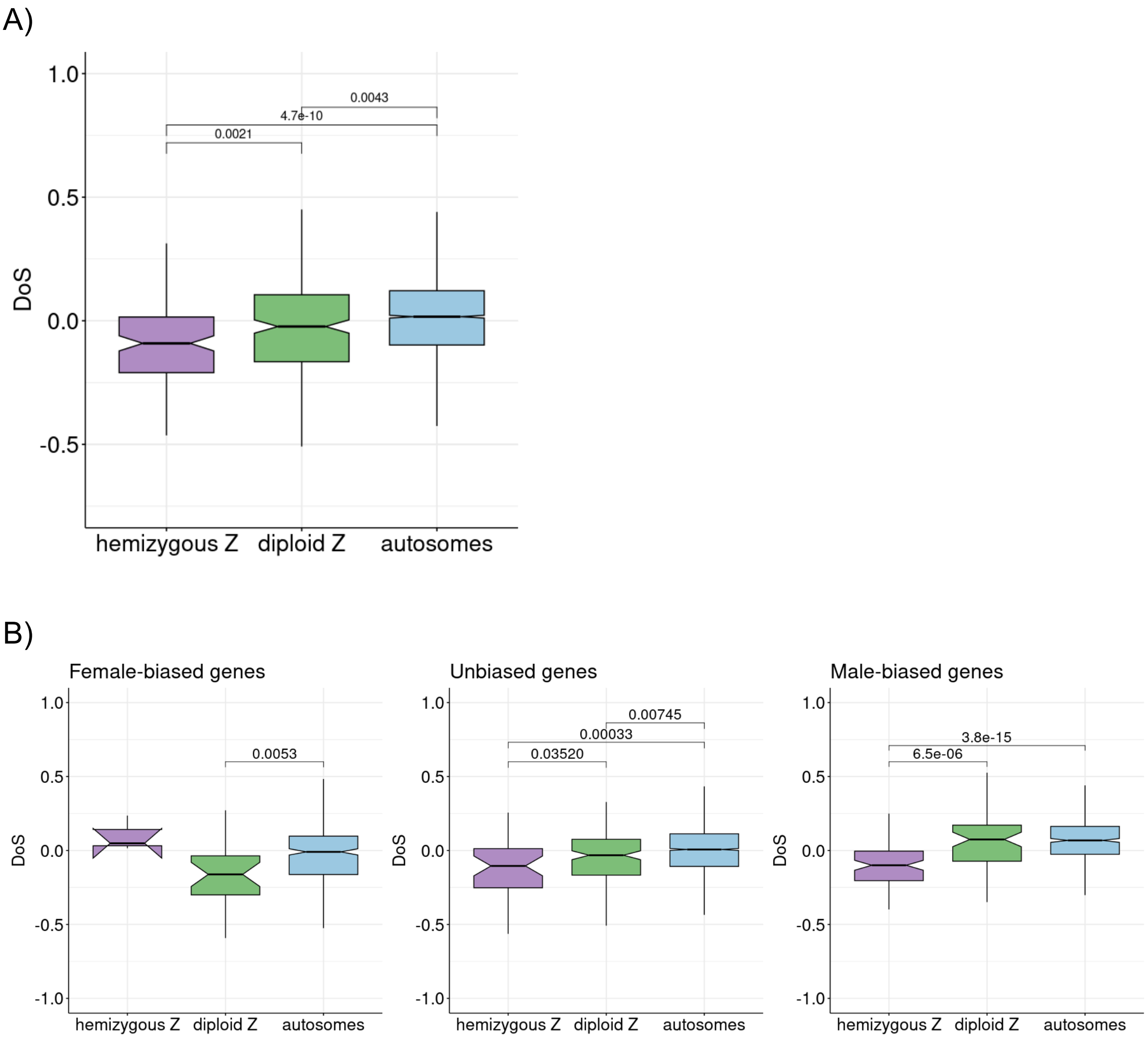


**S.Figure 4**. **A**) Direction of Selection (DoS) as a function of genomic location **B**) Direction of Selection (DoS) as a function of sex-bias and genomic location.

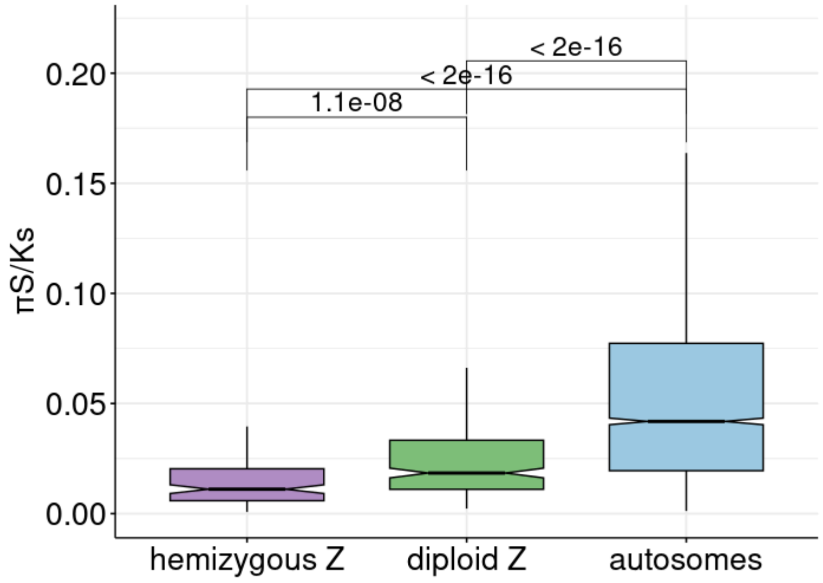


**S.Figure 5.** πs/Ks for hemizygous Z, diploid Z and autosomes
